# Lytic anti-*Staphylococcus aureus* phage M963 resensitizes its host to methicillin by a *tarS*-dependent mechanism

**DOI:** 10.64898/2025.12.19.695355

**Authors:** Marwa Choudhury, Patrick O. Kenney

## Abstract

Bacteriophage M963 targeting a clinical strain of methicillin-resistant *Staphylococcus aureus* (MRSA) was isolated from wastewater. Purification was optimized and the phage was subsequently characterized as a lytic, dsDNA phage within the family *Roundtreeviridae* and genus *Rosenblumvirus*. Host range testing against a small library of *S. aureus* isolates from patients with bacteremia and periprosthetic joint infections showed lytic activity against >60% of tested strains. Evaluation of phage-resistant mutant derivatives of the cognate host showed disruptive mutations within *tarS*, a gene responsible for modifying wall teichoic acid. *tarS* products have been proposed as a target of other members of *Roundtreeviridae.* Synergy testing in MRSA demonstrated phage-antibiotic synergy between M963 and vancomycin, tetracycline, and methicillin. The M963-methicillin synergy prompted testing of the phage-resistant mutant derivatives, showing that phage resistance led to a transient methicillin-susceptible phenotype at clinically relevant minimum inhibitory concentrations in some mutants. Phage M963 is a candidate for compassionate use therapy in appropriate patients.

**Author Summary:** Bacteriophages are viruses that kill bacteria. Because of antibiotic resistance, bacteriophages are being used in the treatment of some infections that are resistant to antibiotics or are not improving despite antibiotics. This manuscript describes the discovery of a bacteriophage, named M963, that kills antibiotic resistant *Staphylococcus aureus* (MRSA). This bacteriophage can kill MRSA from patients with prosthetic joint infections by binding to structures, called wall teichoic acids, on the outside of MRSA. Although the MRSA can evolve to avoid the bacteriophage, this leaves it vulnerable to new antibiotics. By working together with antibiotics, bacteriophage M963 may be a good option for patients with MRSA infections not responding appropriately to standard treatments.

## Introduction

*Staphylococcus aureus* is a major cause of hardware-associated, particularly periprosthetic joint-associated, infections (1). The presence of *S. aureus* alone is an indicator of poor prognosis, with both methicillin-susceptible (MSSA) and methicillin-resistant *S. aureus* demonstrating similarly poor outcomes (2). In one small study of knee periprosthetic joint infections, 53% of patients with retained hardware and *S. aureus* had infection recurrence compared to 10% of patients infected with other organisms (3). Removal of hardware significantly reduced infection recurrence rates in both groups. The reduction in recurrence following hardware removal is likely due to the removal of biofilm as a source of persistent infection (4). Unfortunately, many individuals with hardware infections are not appropriate for removal of hardware for reasons including poor bone stock, comorbidities, and other factors (5–7). Given high rates of recurrence with appropriate medical and maximally safe surgical therapy, alternative approaches are needed. One such alternative approach is bacteriophage (phage) therapy (8). Case reports and case series in a growing body of literature suggest that phage therapy is an appropriate adjunctive approach for some patients (9).

In this report, we describe the isolation of a lytic *Rosenblumvirus* phage targeting a clinical MRSA isolate from a patient with a chronic periprosthetic hip infection. The characterization of this phage offers insights into the clinical application of this and similar phages.

## Results

Phage Isolation. Wastewater samples from multiple municipalities were obtained and tested for the presence of phage as previously described (10). The growth curve of MRSA isolate MRSA1024 in well 3 of a 96-well plate was consistent with the presence of phage, and follow-up plaque assays demonstrated clear plaques with a halo (Figure 1A). The phage was named M963 and carried forward for characterization.

**Figure 1:**
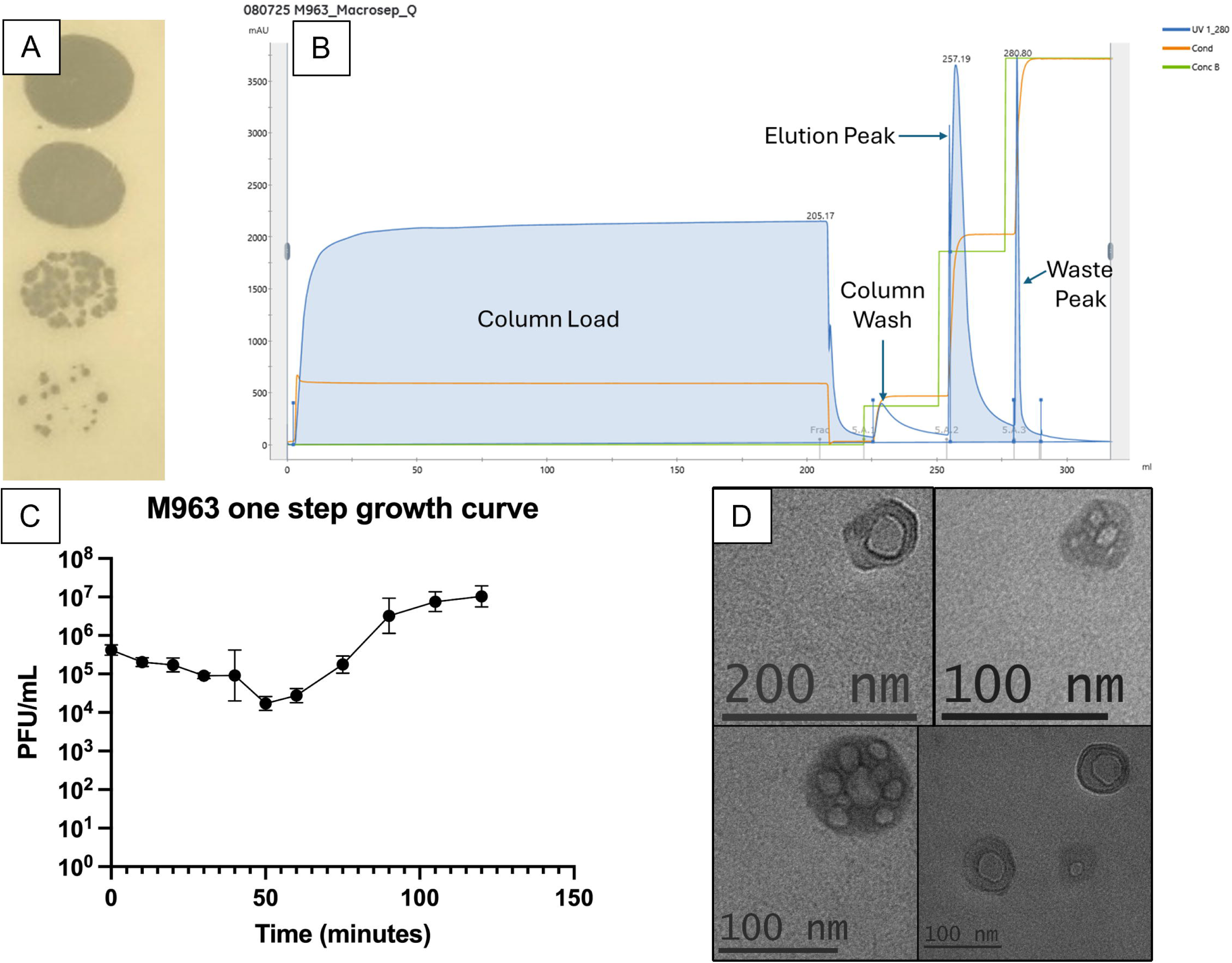
Characterization and purification of phage M963. (A) Serial dilutions of phage M963 on isolate MRSA1024 demonstrate clear plaques or varying size. (B) Sample anion exchange chromatogram for phage M963 with peaks marked. (C) One step growth curve for phage M963 showing a relatively slow lifecycle with a small burst. Error bars represent the mean ± SEM. (D) Multiple representative images from TEM evaluation of phage M963 with scale bars at the bottom left of each panel.

Phage propagation and purification. Phage M963 was propagated against its cognate host, MRSA1024. Phage lysates typically yield counts >10^10^ PFU/mL. Various methods of purification were explored – ammonium sulfate precipitation, size-exclusion chromatography, and cation-exchange chromatography were problematic and had low/minimal yields (data not shown). After multiple pilot studies, the final purification protocol was devised whereby phage lysate was adjusted to a pH ∼10.5 using the addition of 2M Tris base to a final concentration of 0.1M Tris base. This was purified on the Akta Purifier 10 and MacroSep Q resin (YMC) followed by a pH adjusting column wash of 100mM Tris-HCl, 100mM NaCl, pH 8.0, and elution of the major peak with 100mM Tris-HCl, 500mM NaCl, pH 8.0. Concentrations of phage in the eluted peaks were typically near 10^11^ PFU/mL with subsequent buffer exchange and endotoxin removal steps requiring concentration in Amicon 10kDa MWCO filters to achieve a final concentration of 10^10^ PFU/mL. A representative chromatogram is shown in Figure 1B. Subsequent endotoxin testing by gel clot assays showed that the endotoxin levels were <0.25 EU/mL. Sterility of the final vialed formulation was verified by a contract service as per FDA USP-71 guidelines.

One-step kinetics. The basic kinetics of adsorption and burst size of M963 in its cognate host MRSA1024 were explored in triplicate (Figure 1C). Adsorption was slow. A nadir phage concentration was achieved at 40-50 minutes with an adsorption of 96 ± 3%. The latent period was similarly slow, with plateau phage concentrations reached at 105-120 minutes. The burst size was relatively low at 29 ± 49 with a range of 11-50 in individual trials.

Host range. Activity of phage M963 was tested against a panel of 16 total Staphylococcus isolates, including the cognate host MRSA1024 (Table 1). 3 of the 16 isolates were from the same patient, though separated by space and time (MRSA1024, MRSA0925A, MRSA0925B). Other than USA300, included as a reference strain, clinical isolates were utilized from patients with bacteremia and/or bone and joint infections. Phage M963 had measurable, repeatable activity against 11 of the 16 isolates as assessed by either plaque assay or planktonic growth. Plaques were noted on 9 of 16 tested isolates, with efficiency of plating as compared to MRSA1024 shown in Figure 2A. Isolates were also tested in planktonic infection. AUC of control versus phage-treated bacterial isolate were compared, with 6 of 16 isolates demonstrating significant reductions in growth (Figure 2B). Interestingly, 5 isolates with plaque activity did not show reduction in planktonic growth. Conversely, 2 isolates without plaque activity did demonstrate biologically and statistically significant reduction in planktonic growth.

**Figure 2:**
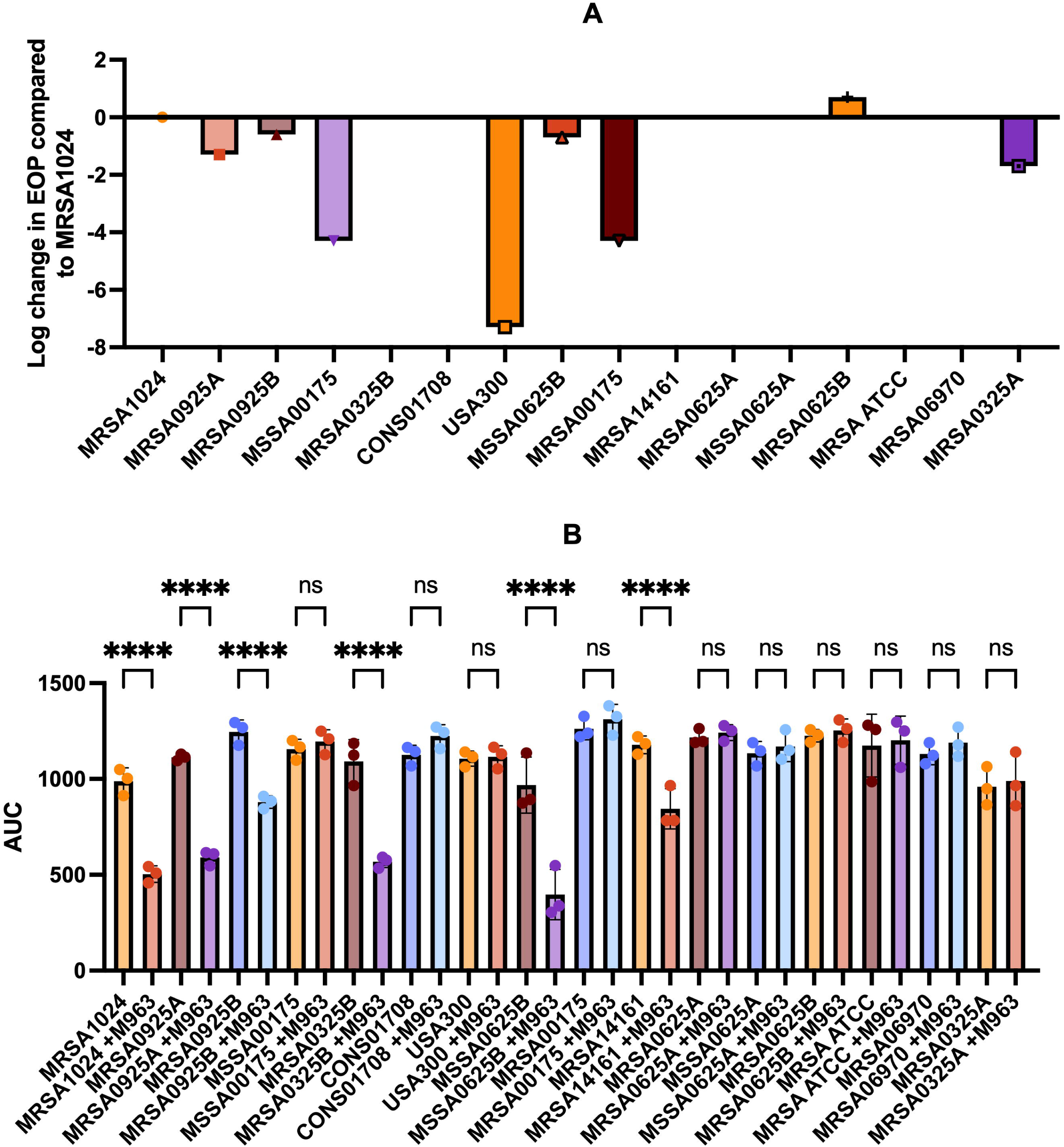
Host range testing for phage M963. The host range of phage M963 was tested by (A) relative efficiency of plating on double agar plaque assays and (B) area under the growth curve in liquid media against a total of 16 *Staphylococcus* strains shown as comparisons of control curves ± M963. Error bars represent mean ± SEM. ns, non-significant. ****, p<0.001.

**Table 1:** Clinical Isolates utilized in testing phage M963 with relevant resistance and source data. NR: Not Recorded.

| Genus | Species | Isolate ID | MRSA (y/n) | Other known resistance | Isolate source |
| --- | --- | --- | --- | --- | --- |
| *Staphylococcus* | *aureus* | MRSA1024 | y | clindamycin, fluoroquinolones | periprosthetic joint infection |
| *Staphylococcus* | *aureus* | MRSA0925A | y | clindamycin, fluoroquinolones | periprosthetic joint infection |
| *Staphylococcus* | *aureus* | MRSA0925B | y | clindamycin, fluoroquinolones | periprosthetic joint infection |
| *Staphylococcus* | *aureus* | MSSA00175 | n | None | bacteremia |
| *Staphylococcus* | *aureus* | MRSA0325B | y | None | periprosthetic joint infection |
| *Staphylococcus* | *epidermidis* | CONS01708 | n | N/A | NR |
| *Staphylococcus* | *aureus* | USA300 (AH-LAC) | y | None | necrotizing fasciitis |
| *Staphylococcus* | *aureus* | MSSA0625 | n | None | periprosthetic joint infection |
| *Staphylococcus* | *aureus* | MRSA00175 | y | None | bacteremia |
| *Staphylococcus* | *aureus* | MRSA14161 | y | None | NR |
| *Staphylococcus* | *aureus* | MRSA0625A | y | None | periprosthetic joint infection |
| *Staphylococcus* | *aureus* | MSSA0625 | n | None | periprosthetic joint infection |
| *Staphylococcus* | *aureus* | MRSA0625B | y | None | periprosthetic joint infection |
| *Staphylococcus* | *aureus* | MRSA ATCC | y | None | bacteremia |
| *Staphylococcus* | *aureus* | MRSA6970 | y | None | NR |
| *Staphylococcus* | *aureus* | MRSA0325A | y | None | periprosthetic joint infection |

Sequencing and bioinformatics analysis. The phage M963 genome is 17,110 bp with 19 genes and no tRNA coding sequences (Figure 3). The annotated genome was submitted under BioProject PRJNA1256892 and can be found on GenBank with accession number PV591857.1.

**Figure 3:**
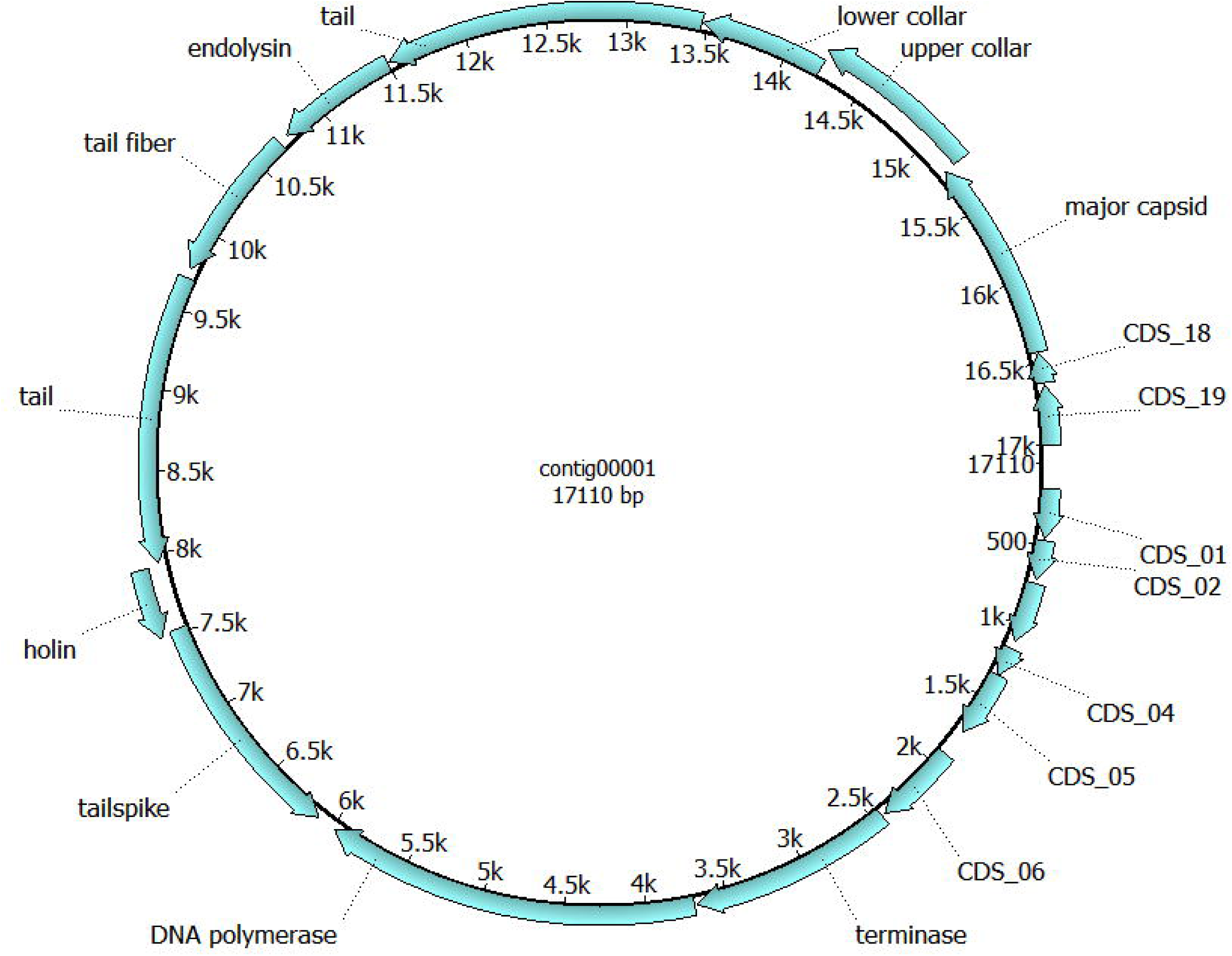
Genome map of phage M963. Phage M963 is represented as a circular genome for convention, with genes marked either with their annotated names or CDS number.

No genes associated with antimicrobial resistance, lysogeny, or other concerning features were noted. The genome is part of a well conserved *Rosenblumvirus* genus. Based on genetic similarity, it likely lies within the same species as a group of phages isolated in Fukuoka prefecture, Japan (accession numbers MK922546.1, MK936475.1, MK922547.1, MK922548.1, MK764384.1, MK903033.1, and NC_055802.1). Further data on these genomes is unpublished at the time of this manuscript. In phage M963, 11 of 19 genes were annotated, with 10 of these genes filling either structural or infection-related roles.

Transmission electron microscopy. A podovirus phenotype was expected based on genome sequencing. Imaging demonstrated an atypical morphology not consistent with the expected phenotype. Multiple representative images shown in Figure 1D. Imaging was repeated with multiple phage lysates and purified formulations, all with similar results. Diameters of the measured particles ranged from 44-91 nm, with a mean of 67 ± 18 nm. Irregular, well-circumscribed inclusions were noted, from one to nine per particle. The inclusions ranged from 5-45nm in diameter without any structural motifs consistent with podoviruses.

Phage resistant bacterial isolates. 4 separate phage-resistant isolates were generated and sequenced in conjunction with the wild type isolate MRSA1024. Mutational analysis performed by Plasmidsaurus software showed consistent gene disruptions across all four isolates affecting *tarS*, a *lapA* family protein, and the intergenic region directly adjacent to the *lapA* family protein (Table 2). The *tarS* sequence was modified by an SNP (R127H) in clone 1 and an insertion sequence leading to a truncated product in clones 2, 3, and 4. The *tarS* gene product is poly(ribitol-phosphate) beta-N-acetylglucosaminyltransferase, responsible for the terminal N-acetylglucosamine on wall teichoic acid. The *lapA* family protein was modified with the same SNP (K7N) in all clones, with extragenic mutations outside the *lapA* ORF at genome positions 1521710 (A>T), 1521716 (G>T), and 1521742 (C>T). AlphaFold3 was utilized to generate theoretical structures on *tarS* and the *lapA* family protein using sequences from the wild type and phage-resistant isolates with a focus on areas with mutations (Figure 4) (11). Only the *tarS* SNP R127H could be directly compared to the wild type via AlphaFold3. The remaining structures, including structures showing the *tarS*-modifying insertion sequence and the *lapA* family protein SNP K7N, were low confidence and not evaluable.

**Figure 4:**
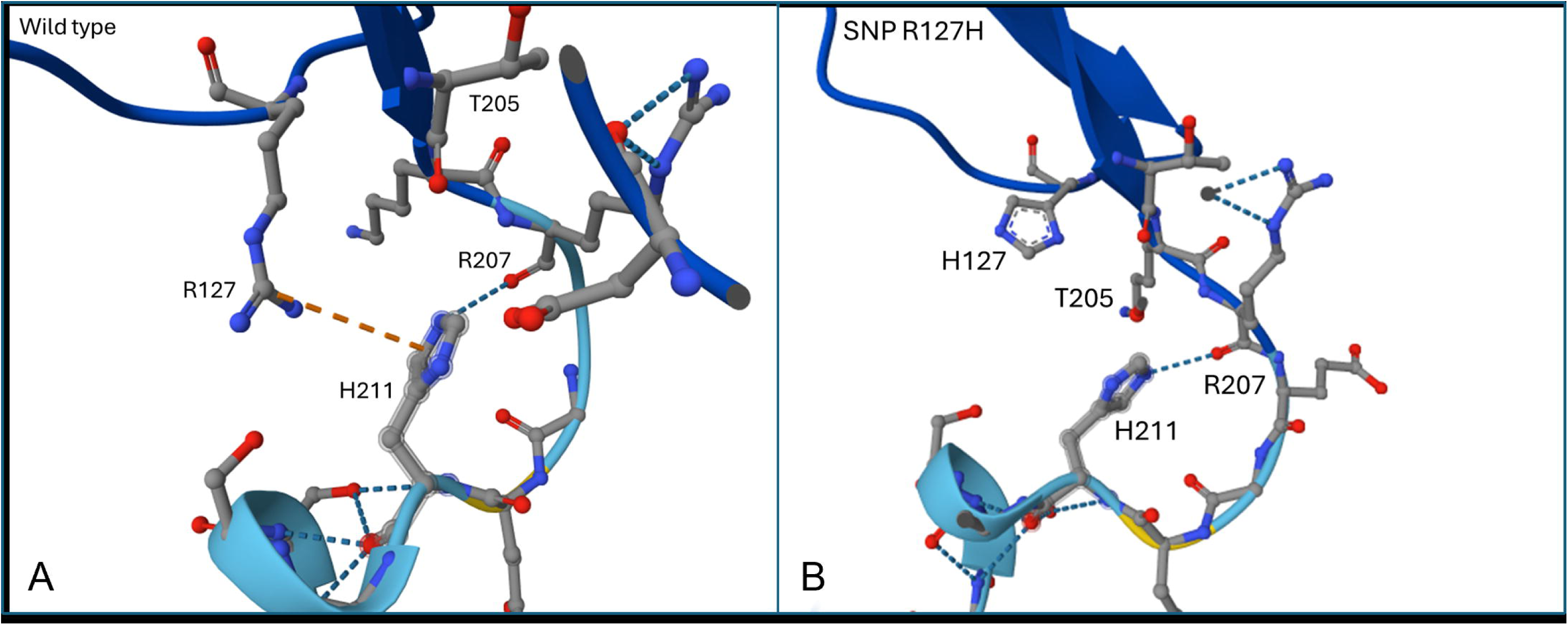
AlphaFold3-generated views of the *tarS* active site. (A) The wild-type MRSA1024 *tarS* active site with relevant amino acids highlighted. R127 is in a cation-pi interaction with H211, which has a hydrogen bond with R207. (B) Phage-resistant mutant with the *tarS* SNP R127H. H127 is rotated away from H211 and non-interactive which T205 rotates to fill the intervening space. H211 and R20 still share a hydrogen bond. As per AlphFold3 notation, very high confidence regions with pIDDT >90 have a dark blue backbone, confident regions (70 < pIDDT < 90) have a light blue backbone, and low confidence regions (50 < pIDDT < 70, only at Glu210) have a yellow backbone.

**Table 2:** Mutations present in phage-resistant strains. Four single colonies of *in vitro*-generated phage M963-resistant MRSA1024 were evaluated by whole genome sequencing. Results are outlined below, with all 4 clones showing modifications in *tarS* and a *lapA* family protein. SNP: single nucleotide polymorphism; VUS: variant of unknown significance.

| Clones | Variant Type | Reference sequence | Start | End | Variant Size | Affected Genes | Mutation |
| --- | --- | --- | --- | --- | --- | --- | --- |
| 1 | snp | Clone 1 | 300669 | 300670 | 0 | *tarS* | R > H |
| 2, 3, 4 | insertion | Clone 2 | 301962 | 303482 | 1520 | *tarS* \| *tnp* | Truncated *tarS*, insertion of ISL3 family IS1181 transposase and sRNA sau-5949 |
| 2, 3, 4 | snp | Clone 2 | 1015528 | 1015529 | 0 |  | Intergenic |
| 1, 2, 3, 4 | snp | Clone 1 | 1521511 | 1521512 | 0 | *lapA* | K > N |
| 1, 2, 3, 4 | snp | Clone 1 | 1521709 | 1521710 | 0 |  | Intergenic |
| 1, 2, 3, 4 | snp | Clone 1 | 1521715 | 1521716 | 0 |  | Intergenic |
| 1, 2, 3, 4 | snp | Clone 1 | 1521741 | 1521742 | 0 |  | Intergenic |

Phage-antibiotic synograms. The ability of phage M963 to complement antibiotic therapy *in vitro* was explored. MRSA1024 was grown in the presence of serially diluted M963 and either vancomycin, tetracycline, daptomycin, or methicillin. Heat maps were prepared demonstrating the ability of these combinations to limit bacterial growth (Figure 5). Synergy scores, a.k.a. the Fractional Inhibitory Concentration Index (FICI), were calculated as previously described (12). The daptomycin-M963 combination was additive (FICI=0.501), while vancomycin (FICI=0.26) and tetracycline (FICI=0.064) were synergistic. Tetracycline synergy was more pronounced at lower antibiotic concentrations. Methicillin was also strongly synergistic despite the methicillin-resistant host isolate (FICI=0.017).

**Figure 5:**
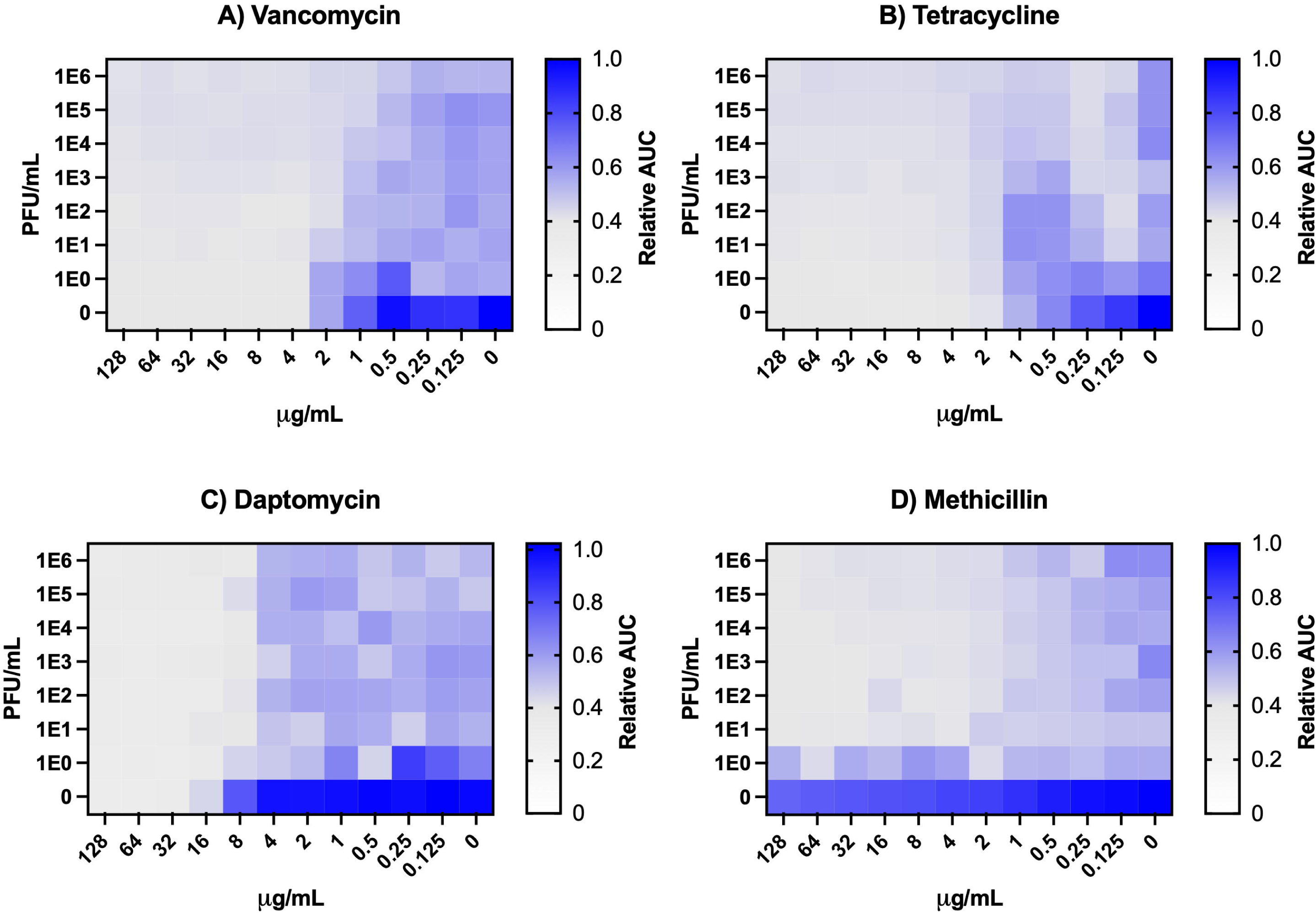
Phage M963-antibiotic synograms in MRSA1024. (A) Vancomycin and M963 have some evidence of synergy as noted at vancomycin concentrations of 1-2 µg/mL. (B) Tetracycline and M963 have notable synergy that improves at the lowest tetracycline concentrations, 0.125-0.25 µg/mL. (C) Daptomycin and M963 demonstrate additive effects without obvious synergy. (D) Methicillin and M963 demonstrate potent synergy at methicillin concentrations ≥2 µg/mL (the CLSI cutoff for susceptible isolates is ≤2 µg/mL).

Methicillin susceptibility by broth microdilution. The phage M963-resistant clones used in sequencing studies above were also tested for methicillin susceptibility in the absence of phage based on the findings in the phage-methicillin synograms (Figure 6). While clones 1, 3, and 4 had an MIC phenotype similar to the wild type isolate when phage was not present, clone 2 was markedly different. In the absence of phage, early growth of MRSA1024 was inhibited by methicillin, but this was not durable, suggestive of a transient phenotypic change.

**Figure 6:**
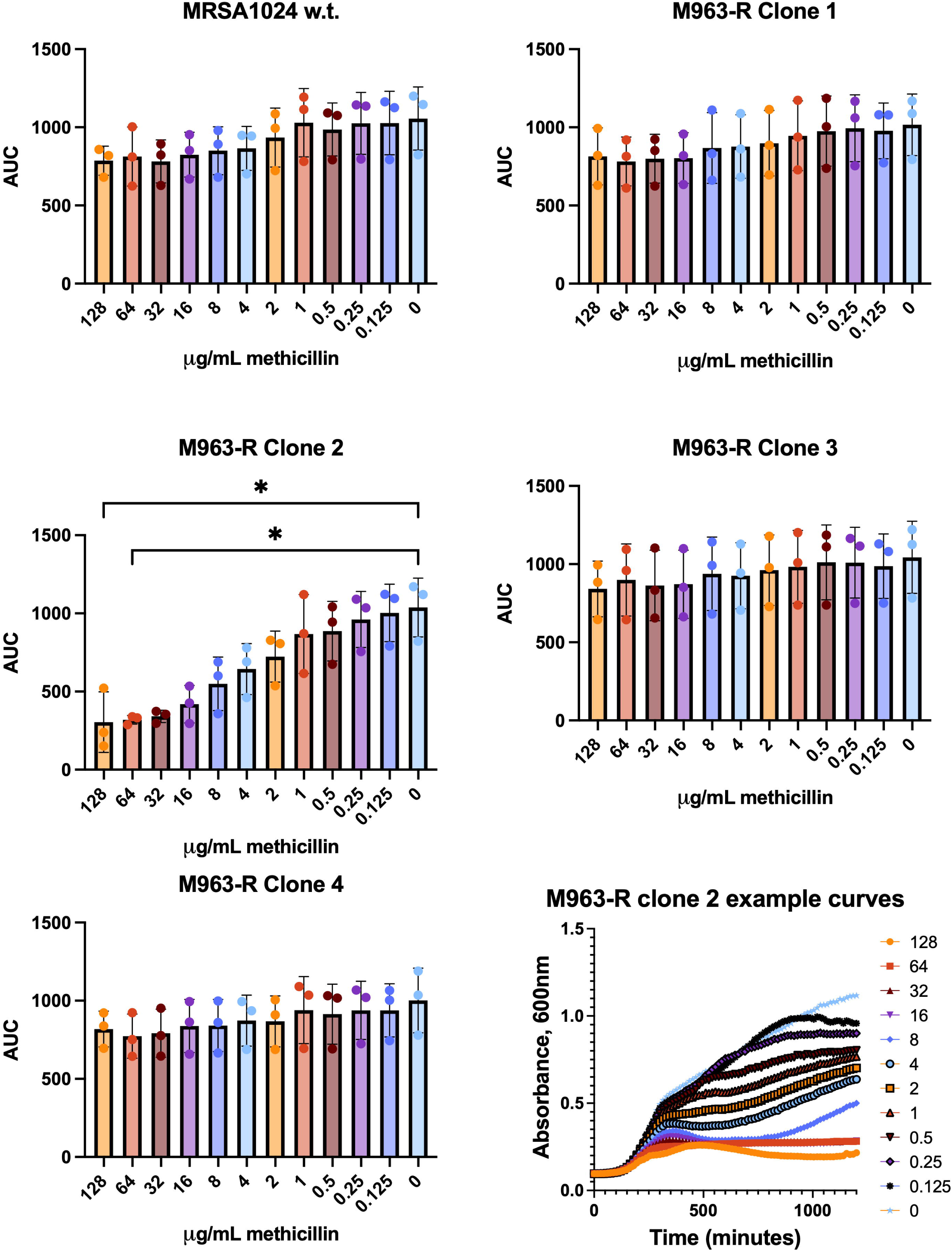
Methicillin testing in MRSA1024 and its phage-resistant mutant derivatives. The areas under the growth curve are shown for varying concentrations of methicillin and (A) w.t. MRSA1024, (B) M963-R clone 1, (C) M963-R clone 2, (D) M963-R clone3, and (E) M963-R clone 4. (F) An example series of curves from one trial for M963 clone 2 demonstrate the presence of methicillin-mediated growth suppression with reversion to methicillin-resistance at earlier timepoints inversely proportional to methicillin concentration for all points ≤8 µg/mL. Error bars represent the mean ± SEM.

## Discussion

Phage M963 targeting the *S. aureus* clinical isolate MRSA1024 was successfully characterized for the purposes of potential phage therapy. While kinetics showed that the phage had a relatively slow life cycle and small burst size on one step growth curves, phage titers >10^10^ PFU/mL were readily achievable. Subsequent purification utilizing anion exchange chromatography yielded a stable, pure phage product in pharmaceutical grade formulation.

Host range testing demonstrated relatively broad activity against a focused library of clinically relevant isolates. 11 of 16 (69%) tested isolates were inhibited by the phage, with 8 of 13 (62%) inhibited once the cognate host and related isolates were removed from the calculation. Phage M963 did not have activity against 3 MRSA isolates, 1 MSSA isolate, and a *S. epidermidis* isolate. This emphasizes a point now well established in the literature – single environmental phages are unlikely to lyse the breadth of a bacterial species (or genus). Larger collections and libraries are required, as well as phage cocktails when used for empiric rather than directed therapy (13, 14).

Sequencing and subsequent bioinformatics analysis showed that phage M963 is part of the well-conserved *Rosenblumvirus* genus of anti-*Staphylococcus* phages. Only 11 of the 19 predicted genes could be annotated.

The TEM findings were unexpected (Figure 1D). The particles noted in TEM were not consistent with podoviruses, though TEM images of multiple close relatives demonstrate a conserved structure (15–17). It is quite possible that the findings are a preparation staining artifact despite multiple imaging attempts.

The mutations found in the phage-resistant clones were generally consistent with one another (Table 2). The first noted mutation is in the wall teichoic acid (WTA) modifying gene *tarS*, whose gene product, poly(ribitol-phosphate)beta-N-acetylglucosaminyltransferase, forms a homotrimer that is responsible for the terminal β-O-GlcNAc in WTA (18). Wall teichoic acids have been well established as phage binding targets in *S. aureus* (19, 20). In the PDBe database, structure 5tzj is a 1.9Å x-ray crystallography structure of *tarS* in complex with its substrate, uridine-diphosphate-N-acetylglucosamine (UDP-GlcNAc) (21). Using comparative sequence analysis, R126 in structure 5tzj is equivalent to R127 in the *tarS* sequence from w.t. MRSA1024. R126 in 5tzj directly interfaces with T204 and the UDP-GlcNAc substrate via hydrogen bonding. AlphaFold-generated structures with pTM scores >0.8 suggest that the SNP R127H in MRSA1024 leads to an altered conformation of the active site (Figure 4). Specifically, an active site-stabilizing R127:H211 cation-pi interaction is lost. The R127H mutation found in phage-resistant clone 1 can therefore be presumed to result in an altered active site. The effect on WTA production was not explored in this study. Sequencing of the remaining phage-resistant clones showed an insertion sequence leading to a truncated *tarS*, with the c-terminal amino acids at 561-573 “VKLNTAHMTYSLK” being replaced by “GSPPNVVGI.” This removes a substantial portion of the terminal beta sheet interfacing with the other two members of the *tarS* homotrimer, theoretically altering the larger structure. Prior studies have demonstrated that only the intact homotrimer maintains enzymatic activity (21). *tarS* is part of a larger system of glycosyltransferases such as *tarM*, *tarO,* and *tarP* (22). The loss of *tarS* function has been shown to result in the use of alternative WTA components (22). This supports our preliminary conclusion that, similar to other anti-*Staphylococcal* phages within *Roundtreeviridae*, the terminal β-O-GlcNAc in WTA is the binding target of phage M963.

The effects of the remaining SNPs found in the sequencing analysis are unclear. The predicted *lapA* family protein is not well studied in *S. aureus*. In *Pseudomonas fluorescens*, a similar protein has been shown to be important in the maintenance of biofilm (23). As this is a simple protein (62 amino acids) with a structure predicted to be composed simply of two sequential alpha helices around a central hairpin turn, the relationship with *lapA* in *P. fluorescens* may be coincidental. AlphaFold3-generated structures of this protein had pTM scores of 0.51 and were not evaluated. The structural effect of the K7N SNP is thus unknown, and the remaining SNPs are in the intergenic region between the predicted lapA family protein and its neighbor, a response regulator transcription factor. The significance of these clustered SNPs warrants further investigation.

*S. aureus*, inclusive of both MSSA and MRSA, is the most common cause of prosthetic joint infections (1). For most infections, current phage therapy approaches involve coadministration of phages and antibiotics (9). A variety of methods have been tested to evaluate the interactions between phages and antibiotics as combinations may be neutral, synergistic, or even antagonistic (12, 24, 25). Synograms, 96-well plates with cross-titered phage and antibiotic combinations, have been used as a static method of evaluating phage and antibiotic interactions *in vitro* (26) and were generated with the permissive phage target MRSA1024, phage M963, and the antibiotics vancomycin, tetracycline, daptomycin, and methicillin (Figure 5). Plates were evaluated both visually and with FICI scores. Daptomycin appeared to have additive effects with phage M963 but no obvious antagonism or synergism. In contrast, vancomycin and tetracycline showed clear synergism with phage M963, particularly with tetracycline at sub-MIC concentrations. Methicillin also demonstrated synergy with phage M963 despite an MIC >128µg/mL in the wild-type isolate. Importantly, mutations in *tarS* have previously been shown to directly impact methicillin resistance (18). Subsequent testing of the phage-resistant clones 1-4 showed that, without cotreatment with phage M963, clone 2 had transient methicillin susceptibility as low as 4µg/mL (Figure 6). This was not durable at a clinically relevant MIC, but together the synergy testing and evaluation of phage resistant mutants clearly demonstrate that concurrent and prior phage treatment can impact antimicrobial resistance phenotypes *in vitro*. Further studies should focus on optimizing and enhancing this effect as well as determining its utility in *in vivo* models.

In this report, we isolated and characterized the lytic phage M963 within the *Rosenblumvirus* genus. This phage was isolated against a MRSA strain from a periprosthetic joint infection and demonstrated plaque formation or growth suppression against 8 of 13 unrelated clinical strains tested. Sequencing of phage resistant mutants confirmed the likely binding site, the terminal β-O-GlcNAc in WTA. Mutations within and near a *lapA* family protein bear further investigation as their impact is unclear and this protein does not appear to be well characterized in gram positive pathogens. Further studies investigating the impact on WTA are also warranted given the finding of potent synergy between M963 and methicillin. Finally, phage M963 is a good compassionate use therapy candidate for difficult-to-treat MRSA infections.

## Material and Methods

### Bacterial strains

The acquisition of clinical bacterial isolates from Kaleida Health Clinical Labs (Buffalo, NY) was approved by the University at Buffalo Institutional Review Board (IRB) under STUDY00005706 contingent upon deidentification. All isolates (listed in Table 1) were stored as glycerol stocks at −80°C prior to use.

### Phage isolation

All work with live phage and bacteria was performed in a BSL-2 laboratory under appropriate safety protocols. Wastewater from a variety of sources was screened against the isolate of interest, MRSA1024, obtained from a patient with chronic hip PJI. Dilute influent from a local plant demonstrated plaques after microplate enrichment as previously described (10). The phage was subcloned x3 by serial plaque picking.

### Phage production, purification, and quality control

An overnight culture of *Staphylococcus aureus* MRSA1024 was diluted 1:100 in LB and phage was typically at a low MOI 0.001-0.0001. The culture was grown overnight, then centrifuged and 0.2 µm-filtered. Phage lysates were pH adjusted to >10, then purified on an Akta Pure 10 using MacroSep Q resin (YMC Bioseparations). After load, the column was washed with 0.1M Tris, pH 8.0, and eluted with the addition of 500mM NaCl. Eluted material was buffer-exchanged to phosphate buffered saline (PBS) over PD-10 columns (Bio-RAD), then 0.2 µm-filtered. For endotoxin determination the ToxinSensor™ Gel Clot Endotoxin Assay Kit (GenScript, New Jersey) was used according to the manufacturer’s instructions with appropriate positive and negative controls. USP-71 sterility testing was performed by Accugen Labs (Addison, IL).

### Plaque assays

LB agar plates were covered with 4 mL molten 0.7% LB agar and 500 µL overnight liquid culture of the bacterial host under evaluation. 4-5 µL aliquots of serial 10-fold phage dilutions were applied in a grid pattern and allowed to dry. Plates were inverted and incubated overnight at 37°C. Plaques were enumerated after 16-20 hours of growth.

### Phage kinetics

One step growth curve: an overnight culture of *S. aureus* MRSA1024 was diluted 1:100 in LB; phage M963 was added to a multiplicity of infection (MOI) of <0.1. Filtered samples were removed every 10-15 minutes and enumerated by plaque assay. Experiments were run in triplicate.

### Host range testing, planktonic growth

Bacterial isolates were grown overnight, then diluted 1:100 into 96-well plates comparing bacterial growth with or without phage at an MOI of 0.1 by tracking OD600 every 15 minutes on the Biotek Epoch2 reader. The area under the bacterial growth curve (AUC) was compared for control versus phage exposed curves. All samples were run with both technical and biological replicates in triplicate.

### Phage genome extraction, sequencing, annotation, and analysis

DNA was extracted using the QIAmp DNA Mini Kit (Qiagen) per the manufacturer’s instructions. DNA was quantified on the Denovix DS-11. DNA electrophoresis in 1% agar with Sybr safe stain was used for quality control (Invitrogen, Grand Island, NY). Sequencing was performed by the University at Buffalo Genomics Core using Illumina HiSeq. Bioinformatics was performed on the Phage Galaxy environment (phage.usegalaxy.eu). Shovill was used for assembly, Prokka for basic annotation, and Abricate to evaluate for problematic genes. Genome visualization was performed with Unipro UGENE 51.0. Phage.ai was used to aid in taxonomic classification and virulence evaluation.

### Transmission electron microscopy

Grids (Square Grid, carbon film on copper, 5-6 nm, 300, Electron Microscopy Sciences, Hatfield, PA) were loaded with purified phages (30 second exposure). The grids were then rinsed once with water (deionized, 30 seconds) and stained with 2% uranyl acetate (45 seconds). A Hitachi HT7800 High Resolution 120kV Transmission Electron Microscope was used for image acquisition with image evaluation and annotation in ImageJ.

### Phage resistant isolates

The permissive host, MRSA1024, was grown in the presence of phage at an MOI of 0.001 for 96 hours to generate a resistant pool which was streaked on phage impregnated LB plates. Individual colonies were selected and amplified, again in the presence of phage at MOI 0.001. This was repeated 4 times, and the 4 clones were sent in conjunction with the wild type MRSA1024 to Plasmidsaurus for DNA extraction and sequencing on the Nanopore platform. SNP evaluation was performed by Plasmidsaurus.

### Phage-Antibiotic Synograms

Antibiotic solutions (daptomycin, vancomycin, tetracycline, and methicillin) were prepared separately at a stock concentration of 512 µg/mL in LB. 100 µL of antibiotic stock was added to each well in column 1 of a 96-well plate. This was then serially diluted 1:2 in LB broth with each step from right to left, with a concentration of 0.5 µg/mL in row 11. Column 12 contained no antibiotics. In a separate 96-well plate, all wells were filled with 180 µL LB broth. Phage M963 at 10^8^ PFU/mL was added to row A in 20 µL aliquots, functionally diluting 1:10. The phage was further serially diluted down the plate, leaving row H without any phage. Starting with column 12, 20 µL from each well on the phage plate was transferred to the corresponding well on the antibiotic plate using a multichannel pipette from right to left. A 1:100 dilution of overnight growth of isolate MRSA1024 in liquid culture was then added to each well in 130 µL aliquots, again from right to left, taking care not to contaminate antibiotic-free and phage-free wells with phage or antibiotic, respectively. This process left well H12 as the untreated control. The plate was then covered with a BreatheEasy™ membrane and transferred to the BioSpa system for incubation, tracking OD_600_ in the BioTek Synergy H1 every 30 minutes. Each plate combination was run in triplicate, evaluated in Graphpad Prism 10.1.0, and visualized as a heatmap for analysis.

### Methicillin MIC in phage-resistant clones

The minimal inhibitory concentration (MIC) of methicillin was assessed with minor modification from the CLSI protocol as follows (27). MRSA1024 was grown in LB overnight. Phage-resistant isolates were grown in LB with the addition of phage M963 to 10^7^ PFU/mL. Overnight bacterial growth was adjusted to a concentration of ∼10^5^ CFU/mL in Mueller-Hinton broth. Methicillin was prepared as a 512µg/mL stock solution and used to generate 11 serial 1:2 dilutions from 128 to 0.125µg/mL with a volume of 50µL per well in a 96 well plate; a 12^th^ well was prepared without antibiotic. 150µL of the dilute bacterial culture was added to each well, and the plate was incubated overnight with OD_600_ reads every 15 minutes and intermitted shaking at 37°C in the BioTek Epoch2. Data was evaluated with GraphPad Prism 10.1.0.

### Structural predictions using AlphaFold3

Protein sequences under investigation were uploaded to the AlphaFold server (alphafoldserver.com/) (11). Structures were generated and compared using AlphaFold3 visualization software. Individual residues were scored as “very high confidence,” “confident,” or “low confidence” in the predicted structures by the AlphaFold3 software. Each structure was given an overall predicted template modeling (pTM) score. Structures with a pTM <0.8 were not evaluated in detail due to low confidence.

### Statistical analyses

#### Host range testing against selected *Staphylococcus* strains

All experiments were performed using biological triplicates. Growth curves were evaluated with mean AUC calculated using Graphpad Prism 10.1.0. Differential AUC values were compared. A 90% threshold was used – i.e. any AUC values for phage-treated curves greater than 90% the AUC of the untreated isolate were not considered to be biologically significant. Statistical significance was assessed via ANOVA.

#### Methicillin MIC in phage-resistant clones

The MIC of methicillin was determined in triplicate as noted above. All tests were performed using biological triplicates, and AUC calculated in GraphPad Prism 10.1.0 were averaged. Mean AUC values were compared to the control well, and statistical significance was calculated by nonparametric ANOVA.

## Acknowledgments

This work would not have been possible without the support of Dr. Thomas Russo, MD CM, and Ulrike Carlino-MacDonald.

